# Characterizing pupil dynamics coupling to brain state fluctuation based on lateral hypothalamic activity

**DOI:** 10.1101/2021.09.22.461385

**Authors:** Kengo Takahashi, Filip Sobczak, Patricia Pais-Roldán, Xin Yu

**Affiliations:** High-Field Magnetic Resonance Department, Max Planck Institute for Biological Cybernetics, 72076 Tübingen, Germany; Graduate Training Centre of Neuroscience, International Max Planck Research School (IMPRS), University of Tübingen, 72076 Tübingen, Germany; Medical Imaging Physics, Institute of Neuroscience and Medicine (INM-4), Forschungszentrum Jülich, 52425 Jülich, Germany; Athinoula A. Martinos Center for Biomedical Imaging, Massachusetts General Hospital, Harvard Medical School, Charlestown, MA 02129, USA

## Abstract

Pupil dynamics presents varied correlation features with brain activity under different vigilant levels. The modulation of brain state changes can arise from the lateral hypothalamus (LH), where diverse neuronal cell types contribute to arousal regulation in opposite directions via the anterior cingulate cortex (ACC). However, the relationship of the LH and pupil dynamics has seldom been investigated. Here, we performed local field potential (LFP) recordings at the LH and ACC, and the whole brain fMRI with simultaneous fiber photometry Ca^2+^ recording in the ACC, to evaluate their correlation with brain state-dependent pupil dynamics. Both LFP and functional MRI (fMRI) data showed opposite correlation features to pupil dynamics, demonstrating an LH activity-dependent manner. Our results demonstrate that the correlation of pupil dynamics with ACC LFP and whole-brain fMRI signals depends on LH activity, indicating a role of the latter in brain state regulation.

## Introduction

Brain state fluctuation, as a fundamental feature of animal brain physiology, is related to arousal regulation during wakefulness, sleep, or under anesthesia^1–3^. It can be assessed using eye-tracking (e.g. pupillometry) as an external index in combination with direct brain functional measurements such as electrophysiological recordings or functional MRI (fMRI)^3–6^. For instance, the slow cortical oscillations of the electrophysiological signals are negatively synchronized with pupil size fluctuations during low arousal states in unconscious animals^5^. Besides measuring cortical brain state changes, studies in subcortical areas have revealed the involvement of the lateral hypothalamus (LH) and multiple brainstem nuclei including the locus coeruleus (LC) and periaqueductal grey in the regulation of brain states^2,7–10^. In particular, LC-based noradrenergic regulation of the anterior cingulate cortex (ACC) activity is in tight relationship with the pupil size regulation^7,8,11,12^, presenting a crucial link between the ACC activity and different sleep states and wakefulness^13,14^. It should be noted that the spontaneous change of pupil size and associated brain states are not solely dependent on the noradrenergic pathways; instead, they are likely orchestrated by multiple subcortical structures^12,15,16^. Interestingly, neural spike rates recorded in the ACC present both positive and negative correlations with pupil size change across trials^16^, suggesting more complex subcortical regulation of brain state-dependent pupil dynamics.

The LH is an important hub for brain state regulation^2^ where different neural cell types co-exist, showing varied activities depending on the underlying arousal state^17–20^. Also, electrical activation of the LH directly causes pupil dilation^21^. The LH activity has been reported to modulate LC-mediated arousal state and pupil dynamics^22–24^. There is anatomical evidence showing direct neural projections of arousal-related nuclei from the LH to the ACC^25^. However, the functional interaction between the LH and ACC has not been fully investigated to elucidate their impact on brain state fluctuation and pupil dynamics.

Here, we measured brain state dynamics with pupillometry, electrophysiology in the LH and ACC, and whole-brain fMRI with simultaneous fiber photometry Ca^2+^ recording in the ACC. We first identified a biphasic distribution of the correlation between the local field potential (LFP) and pupil dynamics in both LH and ACC across different trials of recordings. Based on opposite signs of the correlation (positive vs. negative), we discovered two distinct brain states and performed cross-frequency coupling analyses to identify brain state-specific characteristics of LFP signals for the two states. Specifically, the LFP delta (1-4Hz) phase coupled stronger to amplitudes of higher frequency bands at the ACC in trials where LH-LFP and pupil were negatively correlated, suggesting the trails with negative signs as a lower arousal state^26–28^. Meanwhile, similar to the LFP-based pupil dynamic correlation, the ACC neuronal Ca^2+^ also showed an opposite correlation with pupil dynamics across different trials. Interestingly, in trials with the positive correlation between LH fMRI and pupil dynamics, the ACC Ca^2+^ signal showed a primarily negative correlation with pupil dynamics. This observation suggested that similar brain states could be identified in electrophysiology and fMRI measurements based on LH activity. This work revealed unique LH activity patterns with both LFP and fMRI according to its different coupling features with pupil dynamics, which were associated with different brain states.

## Results

### Simultaneous pupillometry and electrophysiology recordings in ACC and LH

LFP was recorded at ACC and LH simultaneously with pupillometry for 15 minutes to study the brain state-dependent pupil dynamics in anesthetized rats (**Fig.1a**). The LFP power spectrum fluctuation was well-aligned with concurrent pupil diameter changes in a representative trial (**Fig. 1b**). Among different trials, varied correlation features, i.e., positive, negative, or no coupling between pupil dynamics and the power of various LFP frequency bands in the LH and ACC were detected, as shown in **Fig 1c and d** (89 trials from 9 animals). The varied correlation patterns were also detected within the same animal (**Supplementary Fig.1**). In particular, among all frequencies, the oscillation of the delta (1-4 Hz) power was most prominent throughout the entire recording period (**Fig. 1b**). Therefore, correlations of delta power with pupil dynamics were further explored to investigate the LFP coupling between LH and ACC.

**Figure 1.**
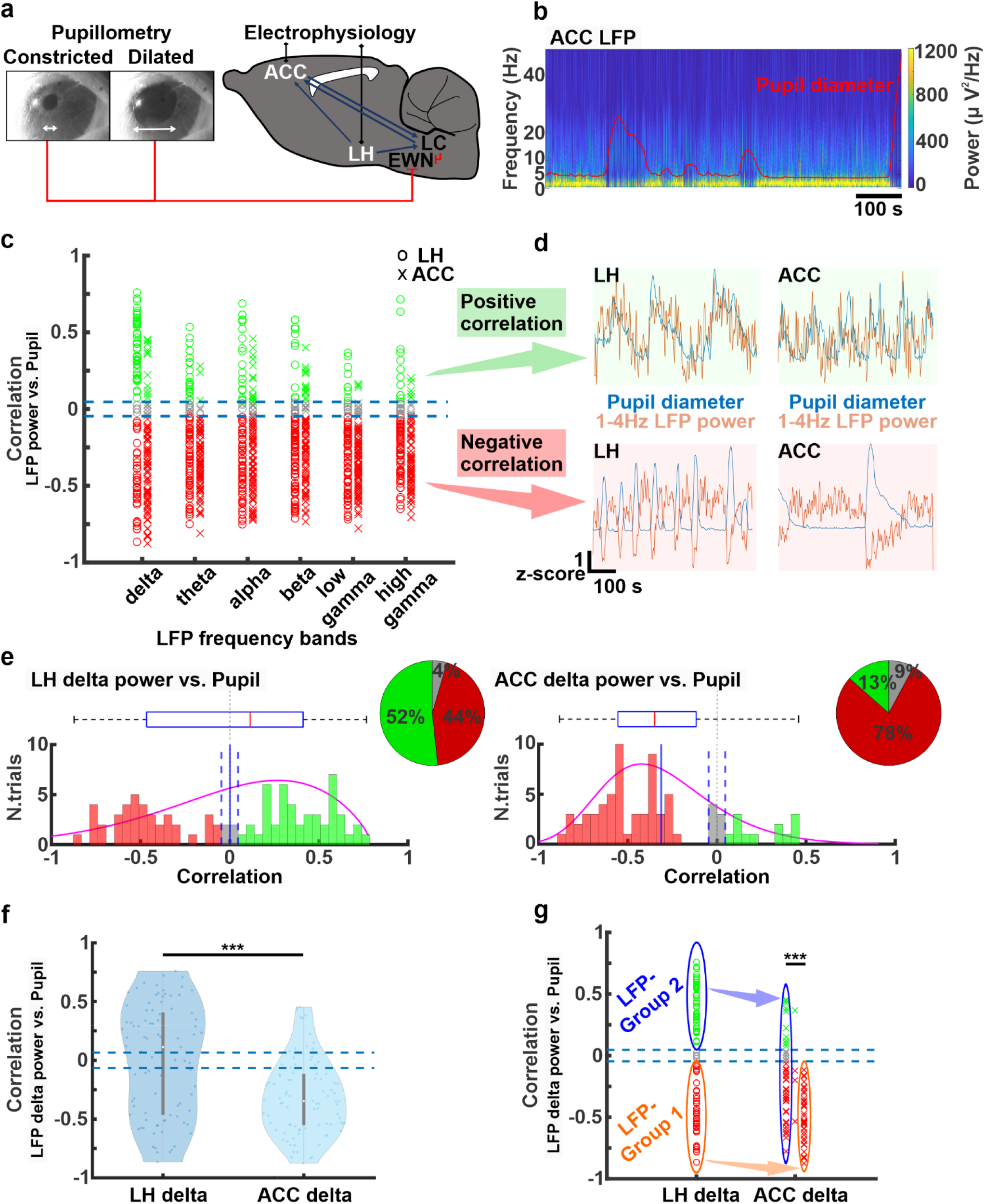
**a)** The experimental setup for simultaneous electrophysiology and pupil size recordings. Images of pupillary response (constricted and dilated pupil) are shown. **b)** Power spectrum of LFP signals in the ACC with pupil diameter fluctuations (red trace) in a representative rat during 15 min of resting state recording. **c)** Correlation coefficients between pupil dynamics and power of all LFP bands in the ACC (circular markers) and in the LH (cross markers) from all trials (n=89, 9 animals). Each marker represents the positive (green), no (gray), or negative (red) correlation on each trial. Thresholds of positive (0.046) or negative (−0.046) correlations are shown as blue dashed lines. **d)** Examples of positive (green shade) and negative (red shade) correlation between pupil dynamics (blue trace) and LFP delta (1-4Hz) band power fluctuations (red trace) in the LH and ACC during 15 min of resting state. **e)** Distribution of correlation coefficients between LFP delta power fluctuation and pupil dynamics in the LH (left; [+ (green)]: n=46, 52%, [0 (grey)]: n=4, 4%, [− (red)]: n=39, 44%) and ACC (right; [+ (green)]: n=12, 13%, [0 (gray)]: n= 7, 8%, [− (red)]: n= 70, 79%). Each pie chart represents ratios of the positive, no, and negative correlations. Vertical blue solid lines represent the mean of correlation coefficients across all trials in the LH and ACC respectively. Pink curves and boxplots represent the distribution of trials for LH and ACC respectively. The vertical black dashed line represents zero. **f)** Violin plots for the distribution of correlation coefficients between pupil dynamics and LFP delta power fluctuation in LH and ACC. **g)** Categorization of correlation coefficients in the ACC between LFP delta power fluctuation and pupil dynamics when LH delta power fluctuation has a positive (LFP-Group 1, orange) or negative (LFP-Group 2, blue) correlation with pupil dynamics. In LFP-Group 1, ACC delta power fluctuation and pupil size fluctuation show a negative correlation (correlation coefficient < −0.046) in all trials. In LFP-Group 2, correlations of ACC delta power fluctuation and pupil dynamics diversify to a great extent. *Note*: [+]: correlation coefficient > 0.046, [0]: −0.046 < correlation coefficient <0.046, [−]: correlation coefficient < −0.046; *** p< 0.001, ** p< 0.01, * p< 0.05.

The correlation between pupil dynamics and LFP delta power fluctuation was more evenly distributed in the LH (positive correlation: 52%, n=46; no correlation: 4%, n=4; negative correlation: 44%, n=39; **Fig. 1e left**) than the ACC, which showed primarily negative correlation to pupil dynamics (positive correlation: 13%, n=12; no correlation: 8%, n=7; negative correlation: 79%, n=70, **Fig. 1e right**). The correlation-based distribution patterns between the LH and ACC differed significantly, as assessed using the two-sample Kolmogorov-Smirnov test^29^ (p = 5.1 x10^−7^; **Fig. 1f**). Based on the positive vs. negative correlation features, we specified two groups, which were statistically determined by calculating the 95% confidence interval from 10,000 correlation measures computed between randomized pupil size and LFP delta power signals (**Supplementary Fig.2**). When the LH delta power and pupil dynamics were negatively correlated (LFP-Group 1), ACC delta power fluctuation and pupil dynamics were negatively correlated in 100% of cases (**Fig.1g**). However, when LH delta power fluctuation and pupil dynamics were positively correlated, possibly indicating a different brain state (LFP-Group 2), the correlation patterns between ACC delta power and pupil dynamics exhibited both positive and negative correlation (**Fig. 1g**). Two-sample t-tests showed that correlations of ACC delta power and pupil dynamics between LFP-Group 1 and 2 occurred differently. (p = 5.2×10^−10^; **Fig. 1g**), indicating distinct brain states based on the relationship between LH and pupil dynamics.

### Characterizing distinct brain states through cross-frequency coupling analyses

To further investigate two brain states identified by distinct coupling features between the LH delta power fluctuation and pupil dynamics (**Fig. 2a**), we performed cross-frequency coupling analyses. In LFP-Group 1, the LH and ACC exhibited the peak power at 2-2.5Hz (**Fig. 2b, Supplementary Fig. 3**). In contrast, in LFP-Group 2 (when the LH delta power fluctuation was positively correlated with pupil dynamics), the LH and ACC exhibited the peak power at 3Hz (**Fig. 2b, Supplementary Fig. 3**). Similarly, analyses of both coherence and phase-locking values between the LH and ACC showed the highest values at 2Hz in LFP-Group 1 but at 3Hz in LFP-Group 2 (**Fig. 2c, d, Supplementary Fig.4, Table.1**). Two-sample t-tests between LFP-Group 1 and 2 phase-locking values showed significant difference at 2Hz (p = 4.8×10^−7^) and 3Hz (p = 8.8×10^−7^). These results indicate that the neuronal coupling between LH and ACC altered between two different states. Moreover, the modulation index, a measurement of the phase-amplitude coupling, between the phase of delta band and amplitudes of all other LFP frequency bands at the ACC was higher in LFP-Group 1 than LFP-Group 2 (**Table. 2**). Two-sample t-tests between LFP-Group 1 and 2 showed significant difference on theta band (p = 1.9×10^−12^), alpha band (p = 4.6×10^−12^), beta band (p = 1.9×10^−11^), low gamma band (p = 1.5×10^−5^), and high gamma band (p = 5.9×10^−8^). These results validate the presence of two distinct brain states involving both LH and ACC under anesthesia, which resembled previous sleep studies^28,30,31^ (see Discussion).

**Figure 2.**
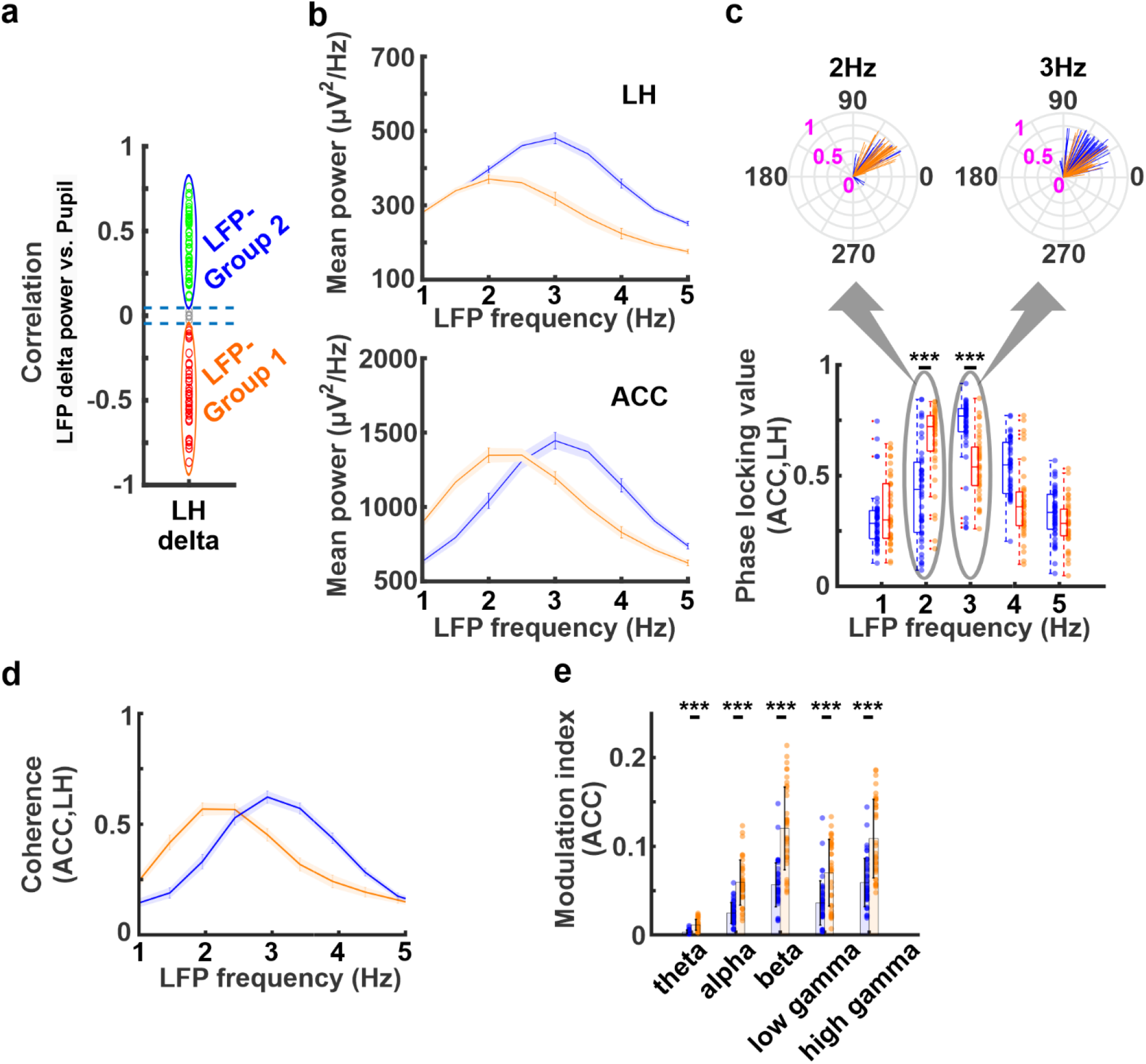
**a)** Correlation coefficients between LFP delta power and pupil dynamics in the LH (green; LFP-Group 2, [+]: n=47,52.8%, grey; [0]: n=3, 3.4%, red; LFP-Group 1 [−]: n=39, 43.8%). **b)** Power spectrum density of the LH (top) and ACC (bottom) in LFP-Group 2 (blue line) and LFP-Group 1 (orange line). The shaded areas represent the standard error of the mean. The peak frequency in both LH and ACC is at 3Hz in LFP-Group 2 (blue line) but at 2Hz in LFP-Group 1 (orange line). **c)** Phase locking value between the LH and ACC in both groups. The highest phase synchrony occurred at 3Hz in LFP-Group 2 (blue) but at 2Hz in LFP-Group 1 (orange). The top radial plots represent the trial-specific phase angles ([degrees], black) and phase-locking values (pink) between LH and ACC at 2Hz and 3Hz. **d)** Magnitude squared coherence between LH and ACC. The highest coherence occurred at 3Hz in LFP-Group 2 (blue) but at 2Hz in LFP-Group 1 (orange). The shaded areas represent the standard error of the mean. **e)** Phase-amplitude coupling using modulation index (MI) at the ACC between the phase of delta (1-4Hz) band and amplitude of other frequency bands. MI is higher in LFP-Group 1 (orange) than LFP-Group 2 (blue) on all frequency bands. Error bars indicate the standard deviation. *Note*: *** p< 0.001, ** p< 0.01, * p< 0.05

**Table 1.**
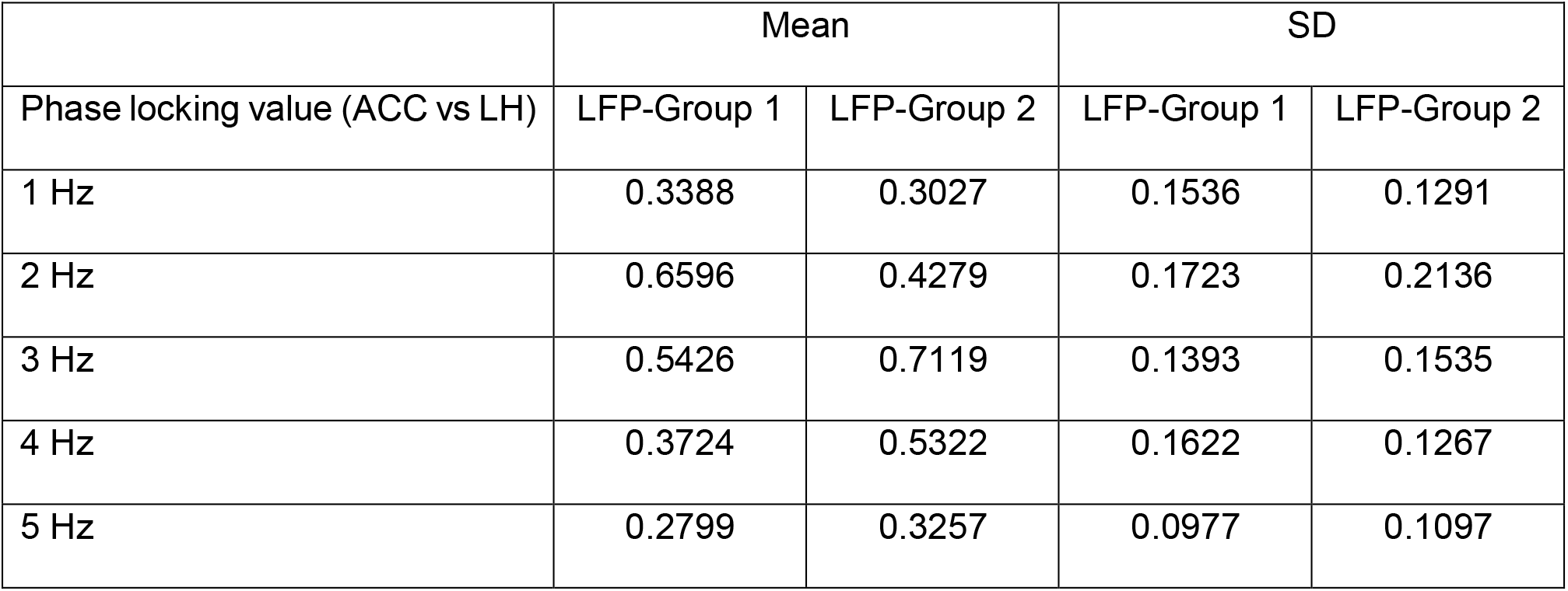
Mean and standard deviation of phase locking value in LFP-Group 1 and 2.

**Table 2.** Mean and standard deviation of modulation index in LFP-Group 1 and 2.

|  | Mean |  | SD |  |
| --- | --- | --- | --- | --- |
|  | LFP-Group 1 | LFP-Group 2 | LFP-Group 1 | LFP-Group 2 |
| Modulation index |  |  |  |  |
| Delta phase vs. theta amplitude | 0.0114 | 0.0035 | 0.0061 | 0.0020 |
| Delta phase vs. alpha amplitude | 0.0590 | 0.0252 | 0.0255 | 0.0122 |
| Delta phase vs. beta amplitude | 0.1196 | 0.0578 | 0.0463 | 0.0263 |
| Delta phase vs. low gamma amplitude | 0.0691 | 0.0376 | 0.0364 | 0.0269 |
| Delta phase vs. high gamma amplitude | 0.1081 | 0.0608 | 0.0439 | 0.0293 |

### Verifying coupling features of neuronal Ca^2+^ signals with simultaneous pupil dynamics and fMRI

To assess the relationship between brain state-dependent pupil dynamics and whole-brain activity, blood oxygen level dependent (BOLD) fMRI with simultaneous pupillometry was performed as previously reported (61 trials from 10 animals)^4^. Meanwhile, we performed fiber photometry to record the GCaMP6-mediated neuronal Ca^2+^ signals in the ACC of these animals, as an alternative to LFP recordings, with better MRI compatibility (**Fig. 3a**)^4^. Similar to LFP results, the power of ACC Ca^2+^ signals also co-varied with spontaneous pupil size fluctuations (**Fig. 3b**). The power of 1-4 Hz Ca^2+^ signal in the ACC was primarily negatively correlated with pupil dynamics (positive correlation: 11%, n=7; no correlation: 20%, n=12; negative correlation: 69%, n= 42; **Fig. 3c, d**), which was consistent with our electrophysiology results (**Fig. 1e**). The clustering of varied correlation features between the 1-4 Hz Ca^2+^ power fluctuation and pupil dynamics among different trials was determined based on the 95% confidence interval calculated by permutation (**Supplementary Fig. 5**). Besides pupil dynamics, ACC neuronal Ca^2+^ signal also showed positive and negative correlation with fMRI signals across different brain regions and trials (**Fig. 3e, Supplementary Fig. 6a**). Interestingly, in trials with a negative correlation between 1-4 Hz Ca^2+^ power fluctuation and pupil dynamics, the Ca^2+^ power fluctuation showed a positive correlation with fMRI fluctuation in cortical and thalamic areas but a negative correlation in subcortical areas, including the LH (**Fig. 3e, f**). In other trials, no significant correlation was detected between fMRI and the ACC Ca^2+^ power fluctuation (**Supplementary Fig. 6b**). These multimodal fMRI results indicated that correlation features between Ca^2+^−based neuronal fluctuation of the ACC and BOLD fMRI signals vary among different brain regions according to different anesthetized brain states.

**Figure 3.**
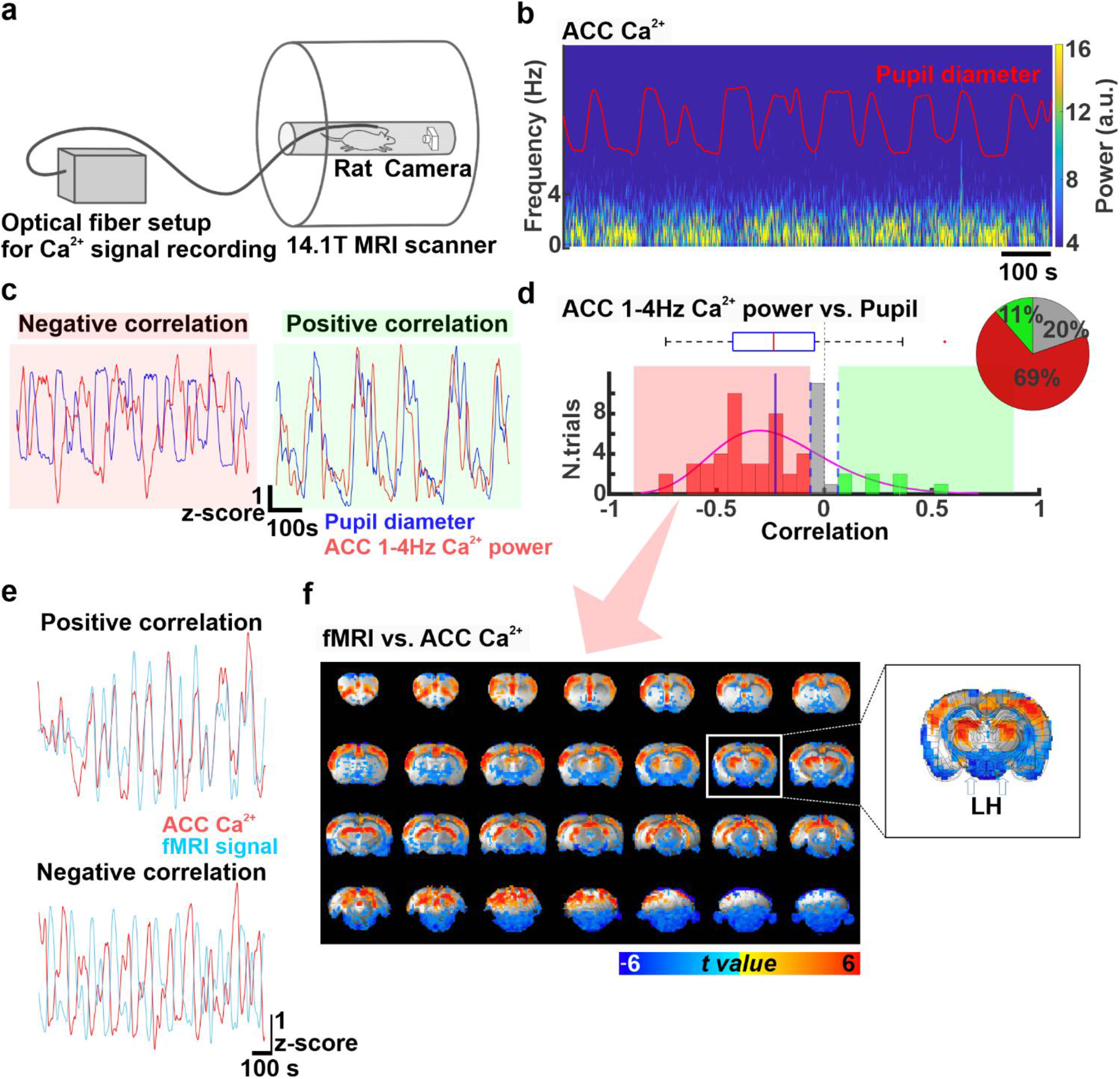
**a)** The experimental setup for multimodal resting-state BOLD fMRI recordings simultaneously with measurements of optical fiber Ca^2+^ signals and pupil dynamics. **b)** Power spectrum of ACC Ca^2+^ signals during 15 min of resting-state along with changes in pupil diameter (red trace) in a representative rat. a.u: arbitrary units. **c)** Examples of positive (right, green) and negative (left, red) correlations between pupil dynamics and ACC 1-4Hz Ca^2+^ power fluctuation during 15 min of resting state. **d)** Distribution of correlation coefficients between pupil dynamics and ACC 1-4Hz Ca^2+^ power fluctuation for all trials (n= 61, 10 animals). These correlation coefficients are negative (correlation coefficient < −0.064) in 69% (n=42, red), none (−0.064 < correlation coefficient < 0.064) in 20% (n =12, grey), and positive (correlation coefficient > 0.064) in 11% (n=7) of the trials. The pie chart represents ratios of the positive, no, and negative correlations. **e)** Examples of positive and negative correlation between fMRI signal (light blue) and ACC 1-4Hz Ca^2+^ power (red) fluctuations during 15 min of resting state. **f)** t-value maps of correlation between ACC 1-4Hz Ca^2+^ power and each fMRI voxel when ACC 1-4Hz Ca^2+^ power fluctuation and pupil dynamics were negatively correlated during 15 min of resting state (n=42,69%). The overlay shows voxels with t-value < −2.25 and t-value > 2.25. A coronal slice at coordinates A/P= −3.2mm is shown magnified on the right, where the location of the LH is indicated with white arrows.

### Classifying distinct brain states based on LH BOLD fMRI and pupil dynamics

Next, we examined if LH BOLD signal could be used to differentiate brain state-dependent pupil dynamics across different trials, as evaluated with LFP data in previous experiments. First, we created a mean correlation map between fMRI and pupil dynamics, showing a global negative correlation pattern as previously reported (**Supplementary Fig. 7**). Moreover, similar to the fMRI-Ca^2+^ correlation, the trial-specific correlation patterns between pupil dynamics and fMRI showed varied signs across different brain regions (**Fig. 4a, Supplementary Fig. 7**). In particular, at the LH, the t-value of correlation coefficient varied extensively across individual trials, showing positive (33%, n=20) or no (67%, n=41) correlations (**Fig. 4b**). We statistically determined this positive correlation by calculating the 95% confidence interval from 10,000 repetitions of correlation measures between randomized pupil signals and each LH fMRI voxel (**Supplementary Fig.8**). In trials showing positive signs, robust positive correlation was also observed between the pupil dynamics and fMRI signal in the brainstem and basal forebrain regions but negative correlation was detected in thalamic and cortical regions (**Fig. 4c**), which was not detected in trials showed no correlation between LH fMRI and pupil dynamics (**Fig. 4c**). Notably, when exploring the correlation of pupil dynamics and ACC 1-4Hz Ca^2+^ power from the trials with the positive LH fMRI-pupil correlation sign, all of the ACC Ca^2+^−pupil correlation coefficients were negative (**Fig. 4d, e;** fMRI-Group 1). This observation was consistent with the electrophysiological results when differentiating LFP-Group 1 and 2 based on LH LFP-pupil relationship (**Fig. 1f**). Two-sample t-tests for correlations of ACC 1-4Hz Ca^2+^ power with pupil dynamics between fMRI-Group 1 and 2 showed a significant difference (p = 3.7×10^−4^; **Fig. 4d**). It should be noted that in LFP-Group 1 the negative LH LFP (delta power)-pupil correlation predicted the negative correlation of ACC LFP (delta power)-pupil correlation, while in the fMRI-Group 1 it was the positive LH-pupil correlation which predicted the negative ACC Ca^2+^-pupil correlation. This might be due to an anti-correlation between BOLD fMRI and delta power in the LH^32^ (see Discussion). In addition, whole-brain fMRI BOLD and ACC Ca^2+^ power coupling was higher in fMRI-Group 1 than fMRI-Group 2 (**Fig. 4f**), indicating distinct brain states associated to the different groups. Overall, these results indicate that correlation of LH fMRI-pupil dynamics can identify distinct brain states, similar to the LH-ACC observations of LFP-based brain states.

**Figure 4.**
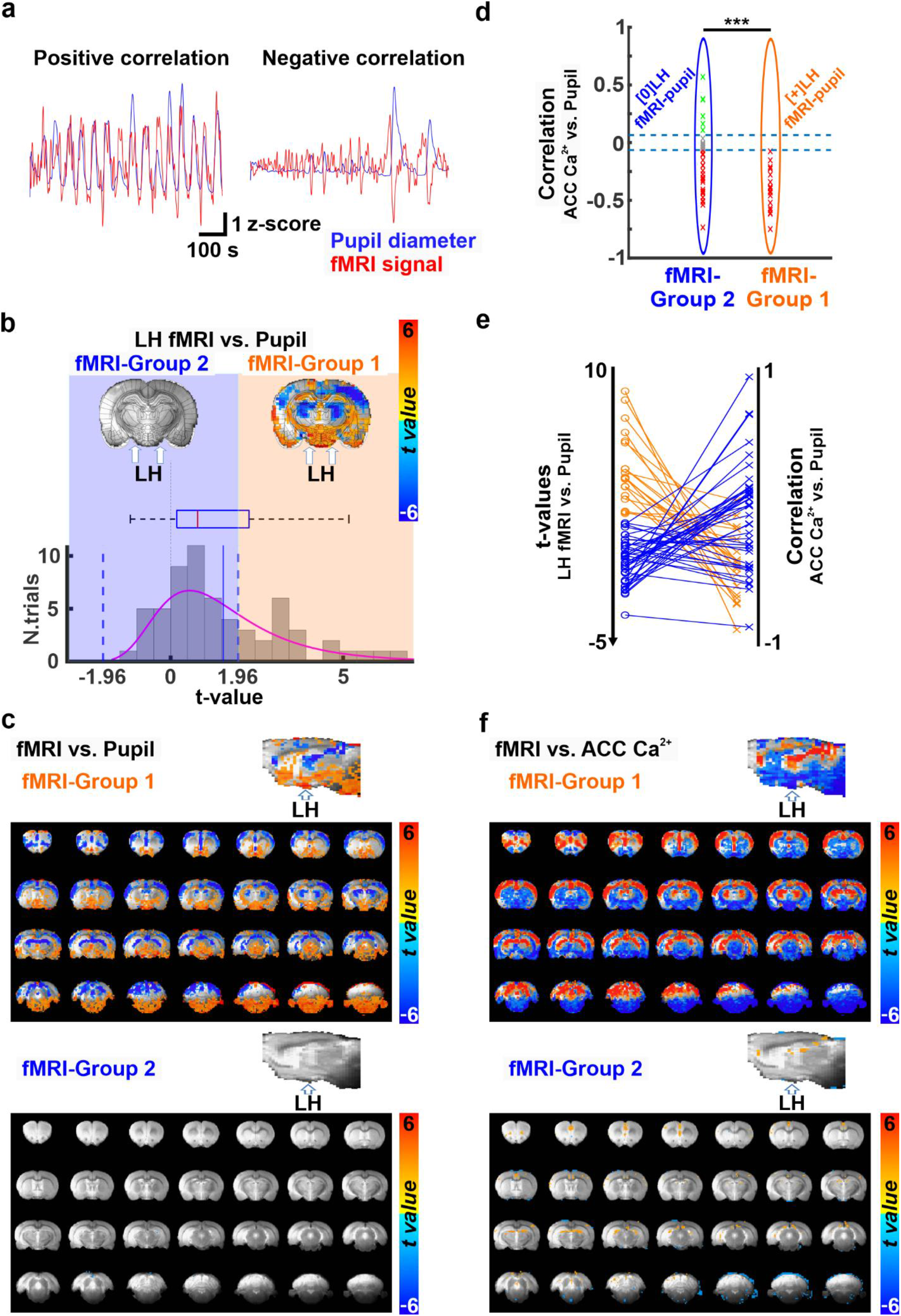
**a)** Examples of positive (left) and negative (right) correlations between pupil dynamics (blue) and fMRI signal fluctuations (red) during 15 min of resting state. **b)** The representation of two distinct brain states. Trials are sorted into two groups characterized by positive (fMRI-Group 1) or no (fMRI-Group 2) correlation between the LH BOLD fMRI signal and the pupil size fluctuations. On the top, two fMRI-pupil correlation maps at A/P= −3.2mm from fMRI-Group 1 and 2 are shown. The location of LH is indicated with a white arrow. The bottom graph represents the t-value distribution of the LH-fMRI and pupil dynamics correlations across all trials (n= 61), on which the state differentiation is based. fMRI-Group 1 (positive correlation) was observed in 33% of the trials (n=20, t > 1.96), and fMRI-Group 2 (no-correlation) in 67% of trials (n=41, t < 1.96). **c)** Whole-brain maps and their sagittal plane view of the correlation between fMRI and pupil dynamics from trials associated with fMRI-Group 1 (top) and 2 (bottom). **d)** Correlation coefficients between pupil dynamics and ACC 1-4Hz Ca^2+^ power fluctuation sorted by fMRI-Group 1 (orange ellipse) and 2 (blue ellipse). The thresholds of correlation coefficients are set at +/− 0.064. **e)** The plot shows the correspondence between two kinds of analysis in the same data: correlation t-values between LH-fMRI and pupil (left) and the correspondent ACC 1-4Hz Ca^2+^ power and pupil dynamic correlation for each trial (right). The color indicates the groups (orange for fMRI-Group 1 and blue for fMRI-Group 2) associated with each trial. **f)** t-value whole-brain maps showing the correlation between fMRI and ACC 1-4Hz Ca^2+^ power from trials assigned to fMRI-Group 1 (top) and 2 (bottom). The overlay shows voxels with t-value < −2.54 and t-value > 2.54. *Note*: *** p< 0.001, ** p< 0.01, * p< 0.05

## Discussion

In the present study, we reported distinct coupling features between brain activity and pupillometry in anesthetized rats, presenting either positive or negative correlation based on multi-modality measurements. When the LH LFP delta power fluctuation was negatively correlated to pupil size changes, the AAC (both LFP and Ca^2+^)-pupil correlation was always negative. In contrast, varied correlation features of ACC-pupil were detected when the LH was positively correlated to pupil dynamics. Based on the LH-pupil relationship, we identified distinct cross-frequency coupling features of the LFP signals detected in the LH and ACC, defining two different anesthetized brain states (LFP-Group 1 vs. 2). In parallel, the trial-specific ACC Ca^2+^-pupil correlation was associated with the fMRI-based LH-pupil relationship similar to the LFP results. In summary, two distinct brain states could be identified based on the LH-pupil relationship, which could be verified by analyzing the site-specific LFP cross-frequency coupling or whole-brain fMRI-based Ca^2+^ and pupil correlation patterns.

We investigated varied coupling features to evaluate different brain states in anesthetized animals. Our results showed two opposite correlation patterns between pupil dynamics and LFP measured at the LH and ACC (**Fig. 1c, d**). It is known that the neural activity in the LH plays an important role in changing and maintaining an ongoing brain state during wakefulness, sleep, and anesthesia^2,20,24,33^. Awake-promoting neurons in the LH (e.g. orexin-producing neurons) reduce their neural firing during both slow-wave sleep^34^ and anesthesia^20^. The anesthetized brain mimics different sleep stages^35^, suggesting common regulatory mechanisms to control the brain state fluctuation. In addition, optogenetic activation of awake promoting neurons in the LH produces cortical arousal and behaviorally awaking from anesthesia^24^ or sleep^17^. Under anesthesia, the neural firing of melanin-concentrating hormone neurons, promoting sleep^19^, are unaffected in spite of the reduced neural firing in awake promoting neurons^20^. Therefore, different neural populations in the LH may contribute to the varied coupling features of pupil dynamics at different arousal states even under anesthesia. Moreover, the LH activation causes pupil dilation^21^ due to inhibition of Edinger Westphal Nucleus (EWN)^9,36^. These attributes of the LH support our observation of two opposite correlation patterns (positive and negative) between the neuronal oscillation in the LH and the pupil dynamics (**Fig. 1c, d**).

We also revealed a unique LH-ACC interaction underlying the varied brain state-dependent pupil dynamics. The ACC is an important hub of the default mode network in both humans and rats^37^, and its involvement in the resting-state connectivity has been observed to differ between high and low arousal states, showing a decreased participation of the ACC in the default mode network during the lower arousal state compared to higher state^38,39^. The neural activity in the ACC has been linked to arousal fluctuation and associated with pupil size changes, which can be explained by its bidirectional connection to the noradrenergic LC^4,13,16,40^. The opposite correlation features between ACC activity and pupil dynamics (**Fig. 1c, d, e**, **Supplementary Fig. 1**) could not be solely explained by noradrenergic projections from the LC. Given the functional and anatomical interaction between LH and ACC^2,13,25,41,42^, our work suggested that LH can play a pivotal role to mediate subcortical regulation of the ACC activity underlying brain state-dependent pupil dynamics. In the state when the LH delta power fluctuation was negatively correlated with pupil dynamics, the ACC LFP-pupil showed negative correlation (**Fig. 1g**). This LFP-based Group 1 state likely corresponds to a lower arousal level based on evidence from the previous electrocorticogram (ECoG) study with pupil size measurements^5^; Yüzgeç et al. (2018)^5^ demonstrated that cortical ECoG-based delta amplitude and pupil dynamics showed very low correlation during wakefulness but strong negative correlation during NREM sleep or moderate negative correlation during REM sleep. We also performed cross-frequency analyses of the LFP signals detected in the ACC and LH, which showed distinct LFP coupling features between LFP-Group 1 and 2 (**Fig. 2b**). Of note, this spectral-based differentiation of brain states is in agreement with sleep EEG studies showing that during NREM sleep, i.e., the deepest state of unconsciousness, the cortical activity peaks around 2Hz^30,31,43^. In contrast, during REM sleep, i.e., a state of relative awareness, the frequency peak of the EEG shifts to higher values (e.g. 2.5-3Hz in frontal-central cortical regions)^31,44^. Besides the frequency-specific coupling features of neural firing in the ACC and LH, the magnitude-squared coherence and phase locking analyses also showed distinct values between LFP-Group 1 and 2, which is similar to previous reports of brain state switches from conscious wakefulness to unconsciousness (e.g. anesthetized or deep sleep state)^45–47^. Since previous studies report that phase-amplitude coupling appears stronger in deeper anesthetized or sleep states^26–28^; our work strongly suggests that the trials in LFP-Group 1 are at a lower arousal level than those in LFP-Group 2.

Pupil dynamics track changes in brain states, in particular, when combined with fMRI as a non-invasive arousal indicator^6,48,49^. It has been shown that pupil dilation occurs concurrently with global negative BOLD-fMRI signal in anesthetized rats^4^, which coincides with decreased low-frequency power recorded in the cortex in both sleeping and anesthetized animals^5,50^. It should be noted that the correlation features between pupil dynamics and fMRI or electrophysiological signals depend on the underlying brain states^4,5,49^. Based on the fMRI signal detected in the LH, we have defined fMRI-Group 1 for positive correlation with pupil dynamics (33% of trials) and fMRI-Group 2 without significant correlation (~67% of trials) (**Fig. 4b**). Our results also showed brain-wide cortical negative fMRI correlation with pupil dynamics in fMRI-Group 1 but the negative correlation was not observed in fMRI-Group 2 (**Fig. 4c**). It is plausible that fMRI-Group 1 corresponds to a lower arousal state than fMRI-Group 2, similar to what has been defined by the LFP-Groups. Indeed, under this positive correlation between LH fMRI and pupil dynamics (fMRI-Group 1), we observed that Ca^2+^-based neuronal signals in the ACC and pupil dynamics were always negatively correlated, a result that is consistent with our electrophysiology findings (**Fig. 4d**, **1g**). Furthermore, our BOLD fMRI and Ca^2+^-based neural correlations were higher in fMRI-Group 1 than in fMRI-Group 2 (**Fig. 4f**), indicating a brain state change based on LH fMRI-pupil relationships.

Although fMRI-pupil interaction shows a promising indicator to estimate brain states noninvasively, more efforts remain needed to better elucidate the coupling feature of fMRI and neuronal signals during brain state fluctuation across different brain regions ^51–54^. In the cortex, He et al. (2008) demonstrated varied correlation between electrophysiological and BOLD signals in awake state and slow wave sleep^54^. Furthermore, depending on eye open/closed arousal states in monkeys, correlation features between fMRI and LFP signals vary^55^, and these eye behaviors can track arousal states with fMRI^6^. The fiber photometry-based Ca^2+^ (GCaMP) recording with simultaneous fMRI^56^ revealed the brain state-dependent correlation features of BOLD signal with neuronal and astrocytic Ca2+ signaling, relying on the neuromodulation from thalamic or brainstem nuclei^4,57^. In our work, ACC 1-4Hz Ca^2+^ power fluctuation was positively correlated with BOLD fMRI signals of the entire cortex (**Fig. 3f**, **Supplementary Fig. 6a**). In contrast, it was negatively correlated with BOLD fMRI in various subcortical regions including the LH, brainstem, and striatum (**Fig. 3f, Supplementary Fig. 6a**). Given the positive coupling feature between LH and ACC LFP data as described in LFP-Group 1, the positive correlation of LH fMRI with pupil dynamics in fMRI-Group 1 suggested negative correlation between LH fMRI and LFP. This negative correlation between BOLD fMRI and LFP low frequency power, in particular at the delta band, has been previously observed in several brain regions including subcortical areas (e.g. striatum)^32,58^, suggesting a non-linear relationship between the neuronal and the vascular network, which is not totally understood.

In summary, we reported varied brain state-dependent coupling features which are orchestrated in multiple regions (e.g. LH and ACC) and cross multiple scales (frequency and amplitude of neuronal firing, inter-region coherence, brain-wide fMRI, pupillary coupling, etc). This work offers a reference framework to understand the LH-ACC relationship, and, more importantly, it presents a novel way to track brain state transitions by bridging pupillary changes with the LH activity. We demonstrate that LH LFP and fMRI signals provide a strong indication of the vigilance level; hence, we anticipate that their assessment may constitute a useful approach to evaluate the ongoing brain states in arousal-related clinical research.

## Method

### Subjects

All experiments were performed in accordance with the European Communities Council Directive (2010/63EU) and the German Animal Protection Law and approved by the Animal Protection Committee of Tübingen (Regierungspräsidium Tübingen, Germany). Nine Sprague Dawley rats (Charles River Laboratories, Sulzfeld, Germany) were used for the multimodal electrophysiology experiments and nine Sprague Dawley rats for the multimodal fMRI experiments. All experiments were performed under alpha-chloralose anesthesia. The multimodal fMRI data had been previously published^4^.

### Surgical procedures for electrophysiology

Prior to the acquisition of the electrophysiology data, rats were anesthetized with 5% isoflurane and orally intubated with a 14-gauge cannula. They were ventilated with a mixture of 2% isoflurane, 30% oxygen, and 70% air at 60±1 breaths per minute by a ventilator (CW-SAR-830/AP) during surgery. The body temperature was maintained at 37-38°C by a heating pad. In order to monitor blood pressure and infuse anesthesia intravenously, cannulation of the femoral artery and vein were peformed using polyethylene tubes (PE-50). Afterwards, rats were placed pronely on a stereotactic device, and two craniotomies were performed using a drill over the ACC and LH. A tungsten electrode (the resistance at 1MΩ; FHC, Inc) was placed at the ACC (M/L= +0.5mm, A/P = +1.2mm, D/V= −1.8mm) and LH (M/L= +1.5mm, A/P = −3.2mm, D/V= −7.8mm) respectively. A bolus of ~80mg/kg alpha-chloralose anesthesia was intravenously administrated into the femoral vein and inhalation of isoflurane was stopped. Afterwards, a mixtures of alpha-chloralose (~25mg/kg/h) and pancuronium (~2mg/kg/h), a paralyzer agent, was continuously infused through an infusion pump during the whole experiment. Once the experiment was done, the location of the electrodes was confirmed by MR imaging.

### Data acquisition of electrophysiology signals

The tungsten electrodes positioned in the ACC and LH respectively were connected to our custom-made pre-amplifier (25x gain and 0.1Hz high pass), and the acquired analog signals were amplified and filtered by MCP-Plus 8 (20x gain and 10kHz low pass; Alpha Omega LTD). The total amplification was therefore 500. A sliver wire was placed between the skull and skin above the cerebellar region as a reference. The analog signals were converted to digital signals by the CED Power1401-3A (Cambridge Electronic Design), and the digital signals were recorded by Spike2 (CED) software at 25 kHz of sampling rate. To ensure synchronization between electrophysiology and pupil size recordings, the start time to record electrophysiology signals was set to be consistent with the start time of pupil size detection by a MacroRecoder software.

### Data acquisition of pupillometry and extraction of pupil diameter

The pupil size was recorded by a customized video camera (dimensions: 30mm × 21mm × 15mm, focal length of the lens: 10mm) in a dark room during the experiments. Infrared LED light (880nm of wavelength) was added in order to acquire a high contrast of the pupil image. The videos were recorded with 30 frames/s, 8 bits per pixel in RGB24, and 628×586 pixels during the electrophysiology experiments and 29.97 frames/s, 8 bits per pixel in RGB24, and 240×352 pixels during the fMRI experiments. The acquired video was processed by the DeepLabCut toolbox^59,60^ in order to extract the pupil diameter from each video frame. Four points were labeled at the edge of pupil, and the pupil diameter was calculated based on these four points with the following formula.

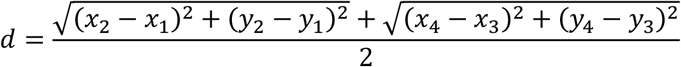

The extracted pupil diameter was first smoothed with a 10 point moving average and then down-sampled to 1 point per 1 second window for fMRI synchronization.

### Data acquisition of MR scans

Prior to the acquisition of MR scans, the same preparatory surgical procedures for electrophysiology such as femoral cannulation were performed. Anesthesia was also provided as described above. All MR scans were performed on a 14.1T / 26cm magnet (Magnex, Oxford) with an Avance III console (Bruker, Ettlingen), as previously described^4^. A custom-made elliptic trans-receiver surface coil (~2×2.7cm) was used for signal transmission and reception. For the acquisition of functional data, a 3D EPI sequence covering the whole brain was used with the following sequence parameters: 1s TR, 12.5ms TE, 48×48×32 matrix size and 400×400×600μm resolution. The total scan time was 15min 25s (925 TRs) for each EPI sequence. For the registration of MR images to a template, an anatomical RARE image was acquired with the following parameters: 4s TR, 9ms TE, 128×128 matrix size, 32 slices, 150μm in-plane resolution, 600μm slice thickness, 8x RARE factor.

### Data acquisition of Ca^2+^ signals

The GCaMP fluorescence signal was acquired from the ACC (M/L= +0.5mm, A/P = +1.2mm, D/V= - 1.8mm) via an optical fiber (FT200-EMT; NA = 0.39; 200μm diameter, Thorlabs), connected to a light path setup, as previously described^4^. The signals were recorded at 5kHz sampling rate using an analog input to digital converter in Biopac 150 system (Biopac, Goleta, CA, USA).

### Analysis of electrophysiology signals

The acquired data were processed in Matlab. In order to extract the power of LFP signals from every second, spectral analysis using wavelet decomposition was performed with 2 seconds sliding windows between 0.5 Hz and 100 Hz. The extracted LFP powers were categorized into different frequency bands based on a previous animal sleep study^5^ as follows: delta (1-4 Hz), theta (4-7 Hz), alpha (7-14 Hz), beta (15-30 Hz), low gamma (30-60 Hz), and high gamma (60-100 Hz). The extracted power time series was first smoothed with a 11 point moving average. The correlation between the power of each frequency band and pupil diameter changes was assessed after synchronizing both signals, using the Matlab corrcoef function.

#### Coherence

Coherence analysis is a standard method to evaluate functional connection of different brain regions in the frequency domain in electrophysiology studies^61^. Prior to evaluating the coherence, the raw data was downsampled from 25000 to 500 sampling points per second. In the coherence analysis between the ACC and LH electrophysiology recordings sites, the Matlab mscohere function was used to calculate the magnitude-squared coherence with a 2 s Hanning window and an overlap of 1s. This function has been used to compute functional connectivity measures in electrophysiology studies^62^.

#### Phase locking value analysis

The phase locking value (PLV) measures synchronization of the phase between two different brain regions to investigate functional connectivity^63^. Prior to calculating the PLV, the raw data was downsampled from 25000 to 5000 sampling points per second. The phase of each LFP was extracted using the Morlet wavelet and the fast Fourier transform from LH and ACC LFP signals respectively. Then, the PLV was calculated with the following formula:

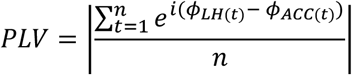

where n represents the number of sampling points, Φ represents phase angles with the same length of sampling points, and *e^i^* represents the exponential integral.

#### Phase-amplitude coupling

The phase amplitude coupling, using the modulation index (MI), is a well-used method to evaluate modulation of brain functions; it estimates how the phase of the low frequency oscillation (e.g. delta band) couples with the amplitude of the high frequency oscillation (e.g. gamma band)^27,64^. Prior to calculating the MI, the amplitude of the upper envelope of the LFP signal in each frequency band was extracted using the Matlab envelope function, and the phase angles of each frequency band were extracted using the Matlab hilbert and angle functions. Then, the MI was calculated using the Kullback–Leibler (KL) distance between the amplitude and phase distributions as described in Tort et al. (2010)^64^ using the ‘modulation_index’ Matlab function (https://github.com/pierremegevand/modulation_index).

### Analysis of fMRI signals

The EPI data from each animal were registered to an anatomical scan (RARE) of the corresponding animal. To spatially match data across animals, one RARE scan was used as a template and other anatomical scans were registered to it, while saving the transformation matrices. Finally, the saved transformation matrices were used to register all functional data together. A bandpass filter was applied to the fMRI signals between 0.01 and 0.1Hz. These fMRI signals were correlated with pupil diameter changes and the power of 1-4Hz Ca^2+^ signals using the Matlab corr function. With the purpose of comparing the whole brain fMRI map with previous publications^4,48^, the voxel-wise correlation t-value was calculated based on the following formula:

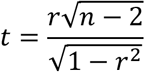

where *r* represents the correlation coefficient and *n* represents the number of sampling points from each run. Once the whole brain t-value map was generated, it was registered on the anatomical image using the AFNI software package^65^. In addition, these anatomical scans were overlapped with a rat brain atlas^66^ to identify the location of brain regions.

### Analysis of Ca^2+^ signals

The acquired optical fiber Ca^2+^ signals (GCaMP) were processed by Matlab. The Ca^2+^ signal was first synchronized with fMRI signals with a customized Matlab code. In order to extract the power of Ca^2+^ signals, spectral analysis using wavelet decomposition was performed with 2 seconds sliding windows between 0.5 Hz and 100 Hz every second. Then, 1-4 Hz of Ca^2+^ powers, equivalent to the delta band of LFP, were extracted. The power of 1-4Hz Ca^2+^ signals was correlated with pupil diameter changes and fMRI signals using the Matlab corrcoef function.

### Analysis to determine the thresholds for the positive, no, and negative correlation

#### ACC and LH electrophysiology vs. pupil

The thresholds for positive and negative correlations were statistically determined by taking the 95% confidence interval from correlations between randomized pupil signals and LFP delta power with 10,000 times of permutation using the Matlab randperm and corrcoef functions (**Supplementary Fig. 2**). Trials above the upper confidence limit (correlation coefficients > 0.065) were defined as positive correlations and those below the lower confidence limit (correlation coefficients < −0.065) as negative correlations. Trials remaining within this 95% confidence interval were considered as uncorrelated. Using the randomized pupil signal, distribution of correlation coefficients with LFP delta power across all trials was examined to assess whether this distribution of correlations with randomized data deviates from the true data. The result shows that distributions between the true and randomized results differ (**Fig. 1e, Supplementary Fig. 2**), indicating that correlation between pupil and LFP signals are not random.

#### Pupil vs. ACC Ca^2+^

The thresholds for positive and negative correlations were statistically determined by taking the 95% confidence interval from correlations between randomized pupil signals and 1-4Hz Ca^2+^ power with 10,000 times of permutation using the Matlab randperm and corrcoef functions (**Supplementary Fig. 5**). Trials above the upper confidence limit (0.065) were defined as positive correlations and below the lower confidence limit (−0.065) were defined as negative correlations. Otherwise, trials within this 95% confidence interval were considered as no correlation. Using the randomized pupil signal, distribution of correlation coefficients with 1-4Hz Ca^2+^ power across all trials was examined to see if this distribution of correlations with randomized data deviates from the true data. The result shows that distributions between the true and randomized results differ (**Fig. 3d, Supplementary Fig. 5**), indicating that correlation between pupil and Ca^2+^ signals are not random.

#### fMRI vs. pupil

The two different brain states (positive or no correlation between BOLD fMRI and pupil dynamics) were statistically determined by taking the 95% confidence interval from correlations between the randomized pupil signal and fMRI signal in the LH with 10,000 times of permutation using the Matlab randperm and corr functions. The result shows that the distribution of correlations with randomized data differs from the true data (**Supplementary Fig. 8**), indicating that correlations between pupil and fMRI signals are not random. Trials above the upper confidence limit (1.96) were defined as positive correlations and those between the upper and lower confidence limit (−1.96 < CI < 1.96) were defined as no correlation. The LH region of interest was located in a single slice centered at the same position as electrophysiology LH recordings (33 manually selected voxels at A/P = −3.2mm). T-values for correlation were calculated for the whole brain analysis as described above. In addition, the correlation between pupil and each fMRI voxel was corrected by the false discovery rate using the ‘FDR’ Matlab function with alpha 0.05 for 20022 voxels (https://www.mathworks.com/matlabcentral/fileexchange/71734-fdr-false-discovery-rate). Thereby, the whole-brain maps show voxels of positive correlation with t-value > 2.54 and negative correlation with t-value < −2.54 (**Fig. 4c, Supplementary Fig. 7**).

#### fMRI vs. ACC Ca^2+^

T-values for correlation were calculated between the power of 1-4Hz ACC Ca^2+^ and each fMRI voxel as described above. In addition, the correlation was corrected by the false discovery rate using the ‘FDR’ Matlab function with alpha 0.05 for 20022 voxels as described above. Thereby, the whole-brain maps show voxels of positive correlation with t-value > 2.25 and negative correlation with t-value < −2.25 (**Fig. 3f, 4f, Supplementary Fig. 6a, 6b**).

## Author contributions

Research design: X.Y., K.T., Data acquisition: K.T., P.P.-R., Data analysis: K.T., Technical support: F.S., Writing – original draft: K.T., Writing – review & editing: X.Y., K.T., P.P.-R., F.S., Supervision: X.Y.

## Acknowledgments

We thank Dr. R. Pohmann, Dr. K. Buckenmaier, Ms. H. Schulz, and Dr. J. Engelmann for technical support, Dr. E. Weiler, Dr. P. Douay, and Ms. R. König for animal protocol and maintenance support, and Mr. M. Arndt and Mr. O. Holder for electrical support.

## Supplementary information

### Characterizing pupil dynamics coupling to brain state fluctuation based on lateral hypothalamic activity

**Supplementary Figure 1.**
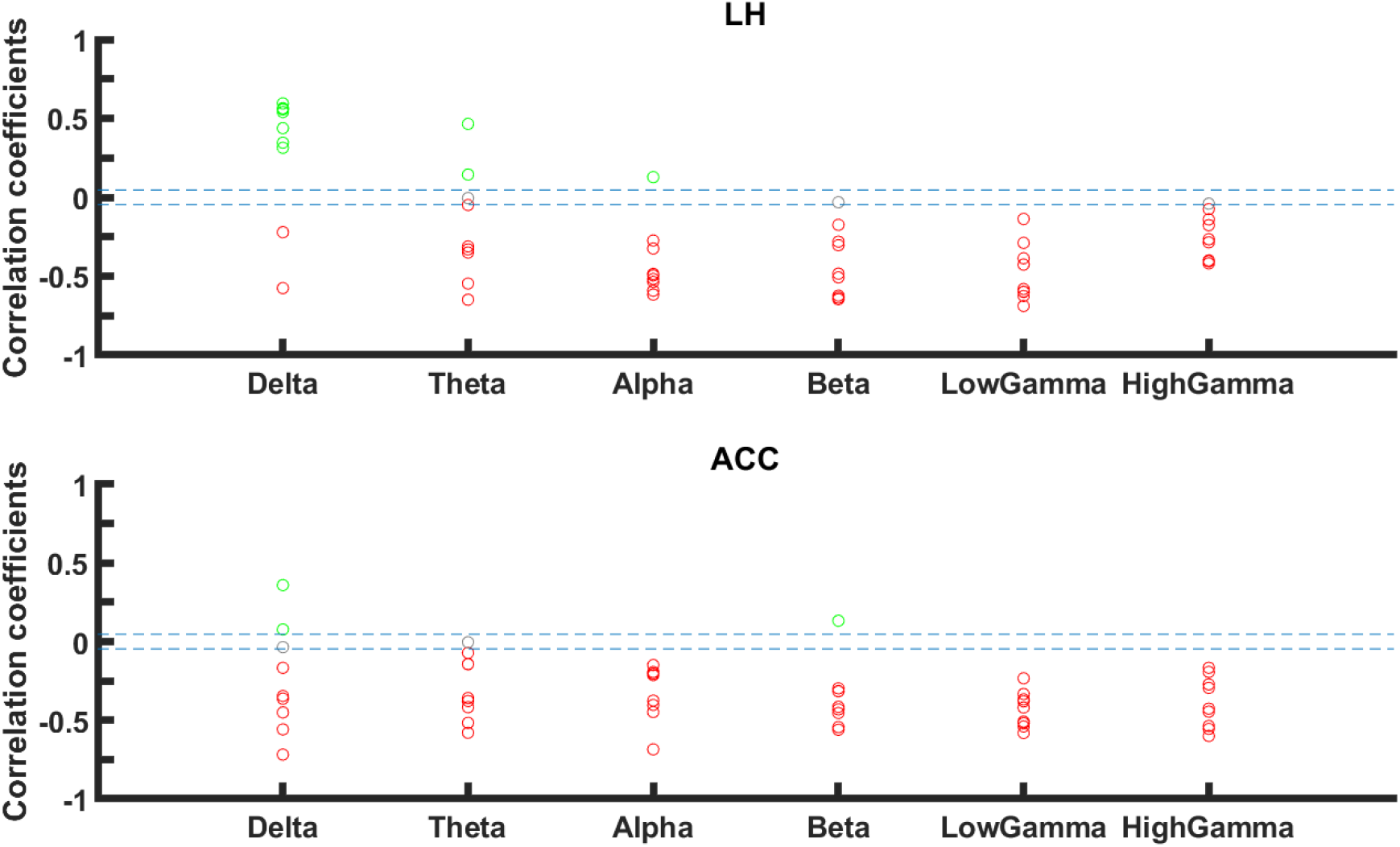
Correlation coefficients between pupil dynamics and power of all LFP bands in LH (top) and ACC (bottom) from a single representative animal. Each marker represents positive (green), no (gray), or negative (red) correlation from every single trial. Thresholds of positive (0.046) or negative (−0.046) correlation are shown as blue dashed lines.

**Supplementary Figure 2.**
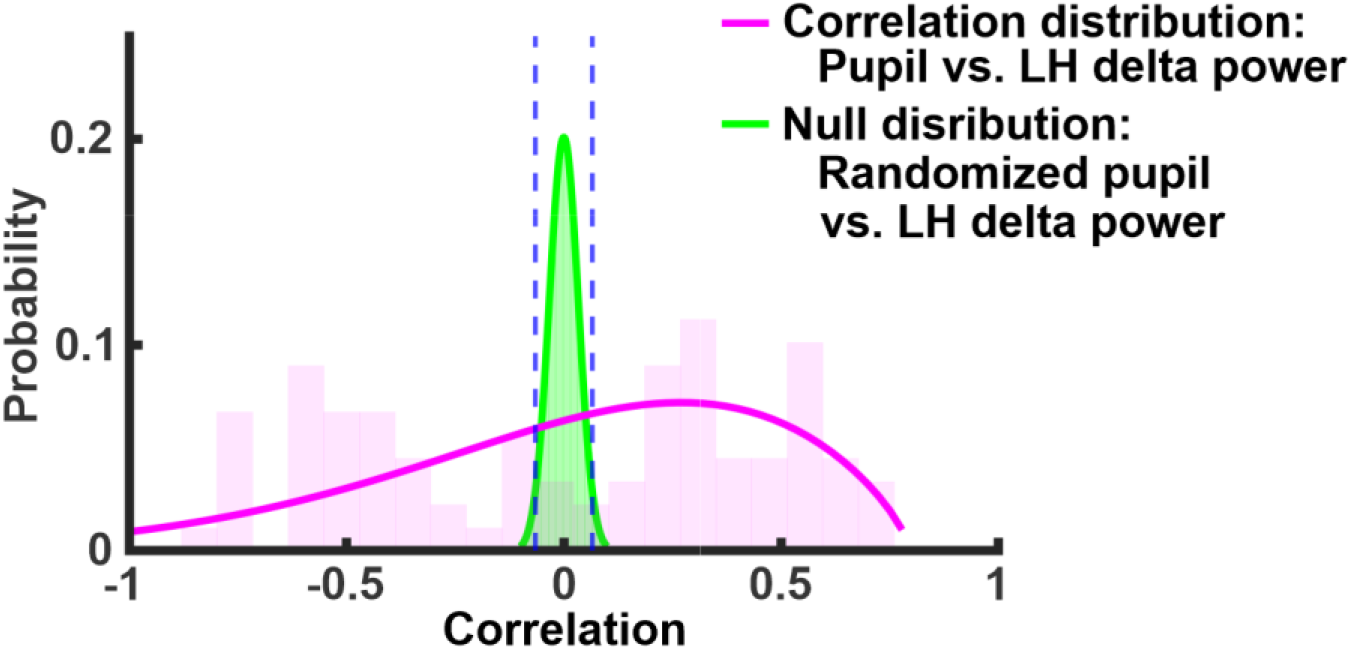
Determination of the threshold for positive and negative correlations between electrophysiology signals and pupil fluctuation. The pupil signals were first randomized using randperm function in Matlab. Using these randomized pupil signals together with 1-4Hz LFP power, the correlation coefficient was calculated, and the permutation was conducted with 10,000 times of repetition across 61 trials. The 95% confidence interval of correlation coefficients from this permutation test was between −0.046 and 0.046, indicated with dashed blue lines. The distribution of correlation coefficients between randomized pupil and LH delta power is in green and the distribution of correlation coefficients between pupil and LH delta power is in pink (same as Fig.1e left).

**Supplementary Figure 3.**
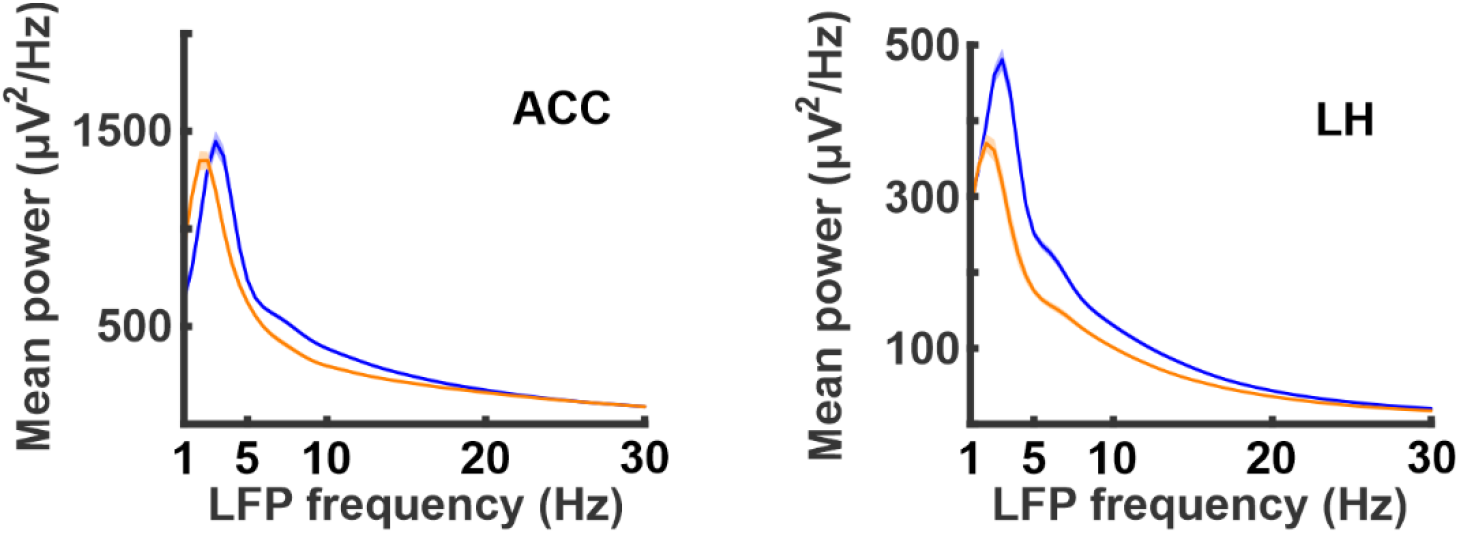
Power spectral density of the ACC (**left**) and LH (**right**) between 1Hz and 30Hz range in [+] pupil×LH LFP(*δ*) cases (blue line) and [−] pupil×LH LFP(*δ*) cases (orange line). Standard errors are represented with a shading area. The peak frequency in both LH and ACC is at 3Hz in LFP-Group 2 (blue; [+] pupil×LH LFP(*δ*) cases) but at 2Hz in LFP-Group 1 (orange; [−] pupil×LH LFP(*δ*) cases).

**Supplementary Figure 4.**
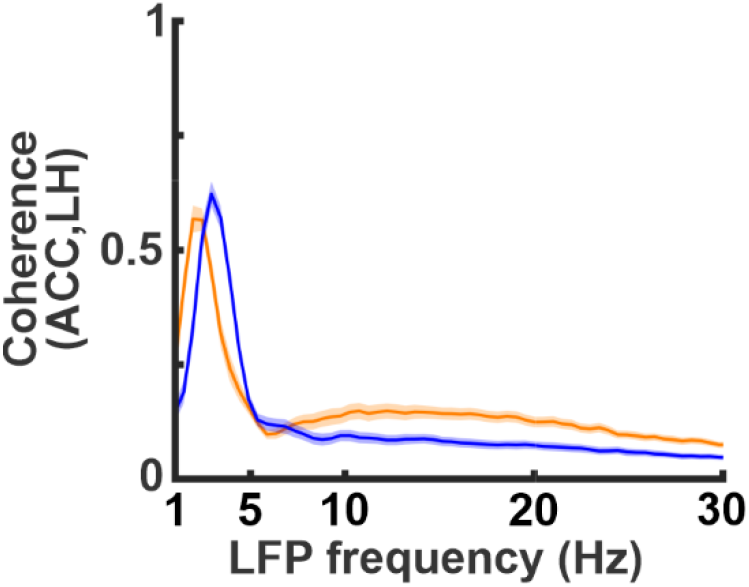
Magnitude squared coherence (MSC) between the LH and ACC in 1Hz to 30Hz range. The highest coherence occurred at 3Hz in LFP-Group 2 (blue; [+] pupil×LH LFP(*δ*) cases) but at 2Hz in LFP-Group 1 (orange; [−] pupil×LH LFP(*δ*) cases).

**Supplementary Figure 5.**
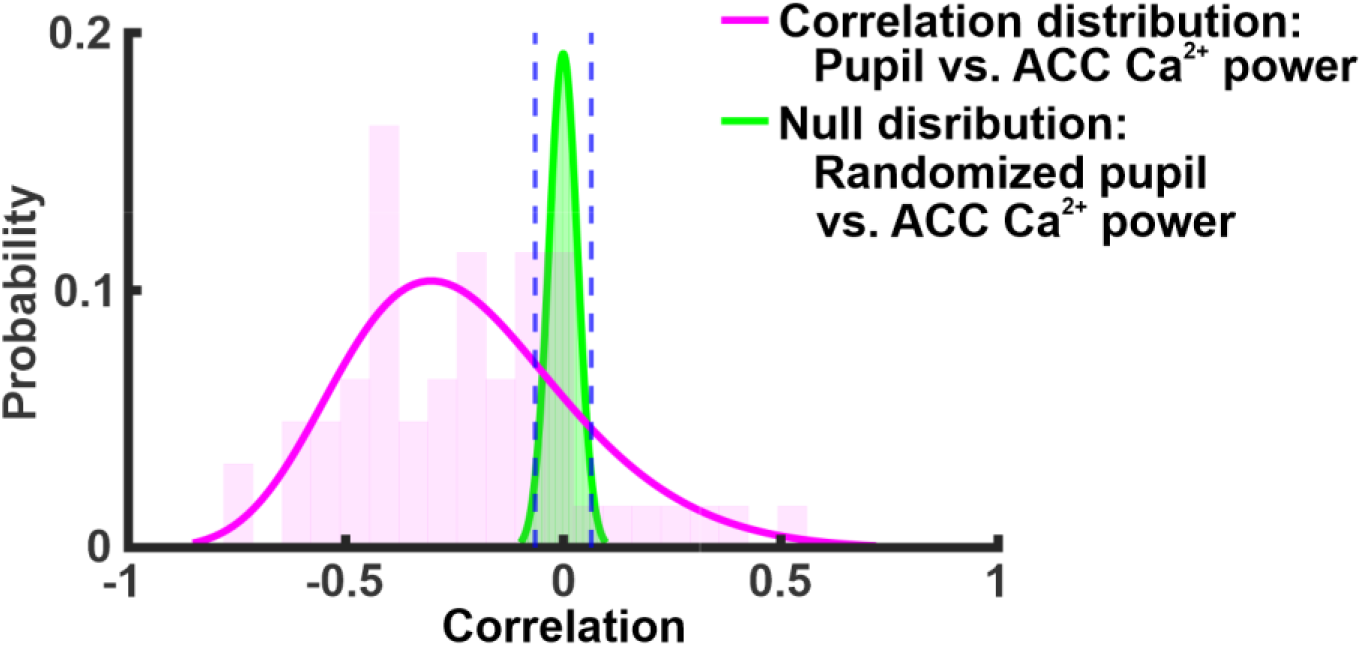
Determination of the threshold for positive and negative correlations between Ca^2+^ signals and pupil fluctuation. The correlation coefficient was calculated between randomized pupil signals and 1-4Hz Ca^2+^ power, and then permutation was conducted with 10,000 times of repetition across 61 trials. The 95% confidence interval of correlation coefficients from this permutation test was between −0.064 and 0.064, indicated with dashed blue lines. The distribution of correlation coefficients between randomized pupil and ACC 1-4Hz Ca^2+^ power is in green and the distribution of correlation coefficients between pupil and ACC 1-4Hz Ca^2+^ power is in pink (same as Fig.3d).

**Supplementary Figure 6.**
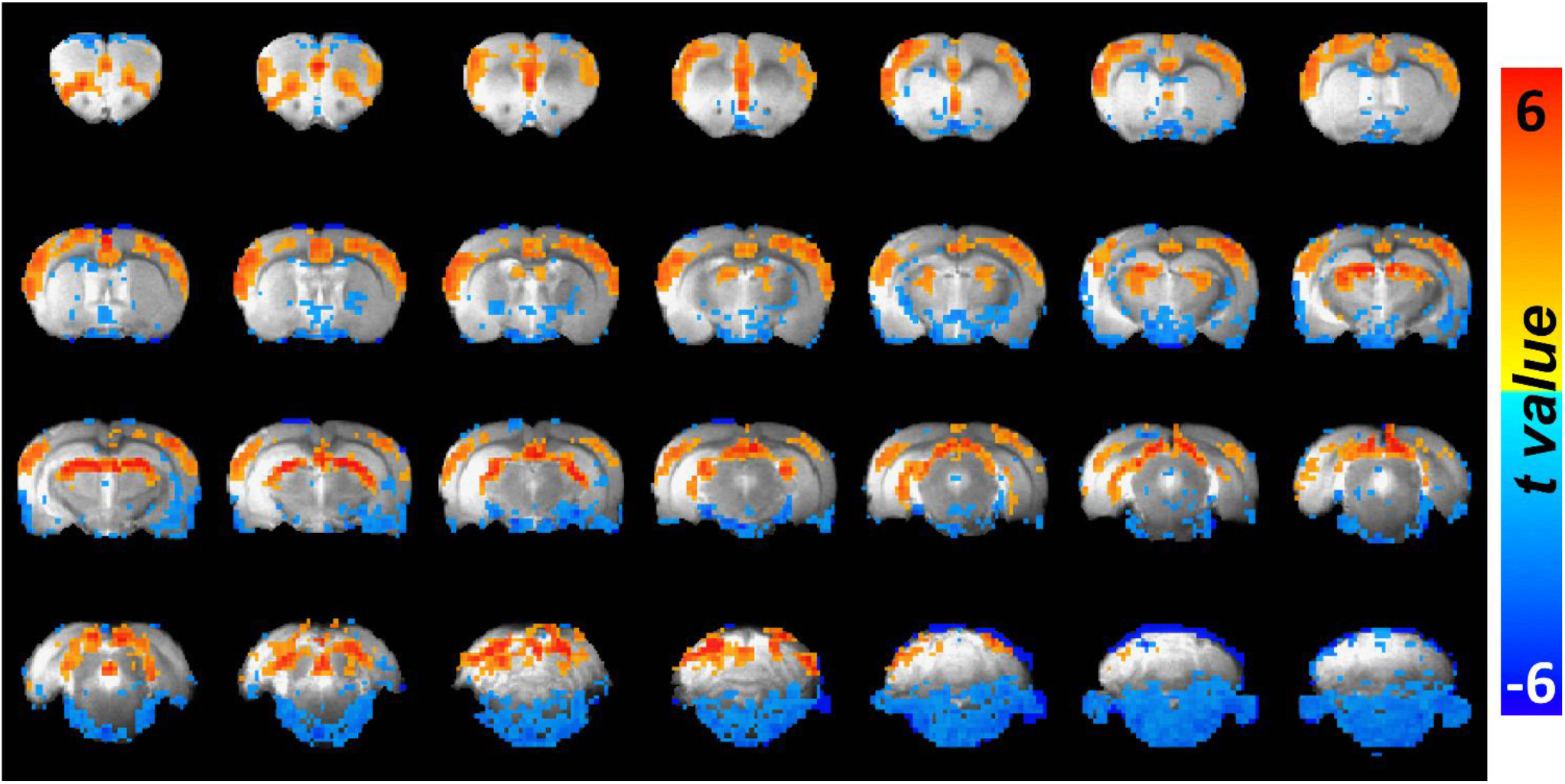
**a)** Averaged correlation maps between ACC 1-4Hz Ca^2+^ power and each voxel of fMRI signals from all trials (n=61). The overlay shows voxels with t-value < −2.25 and t-value > 2.25.

**Supplementary Figure 6.**
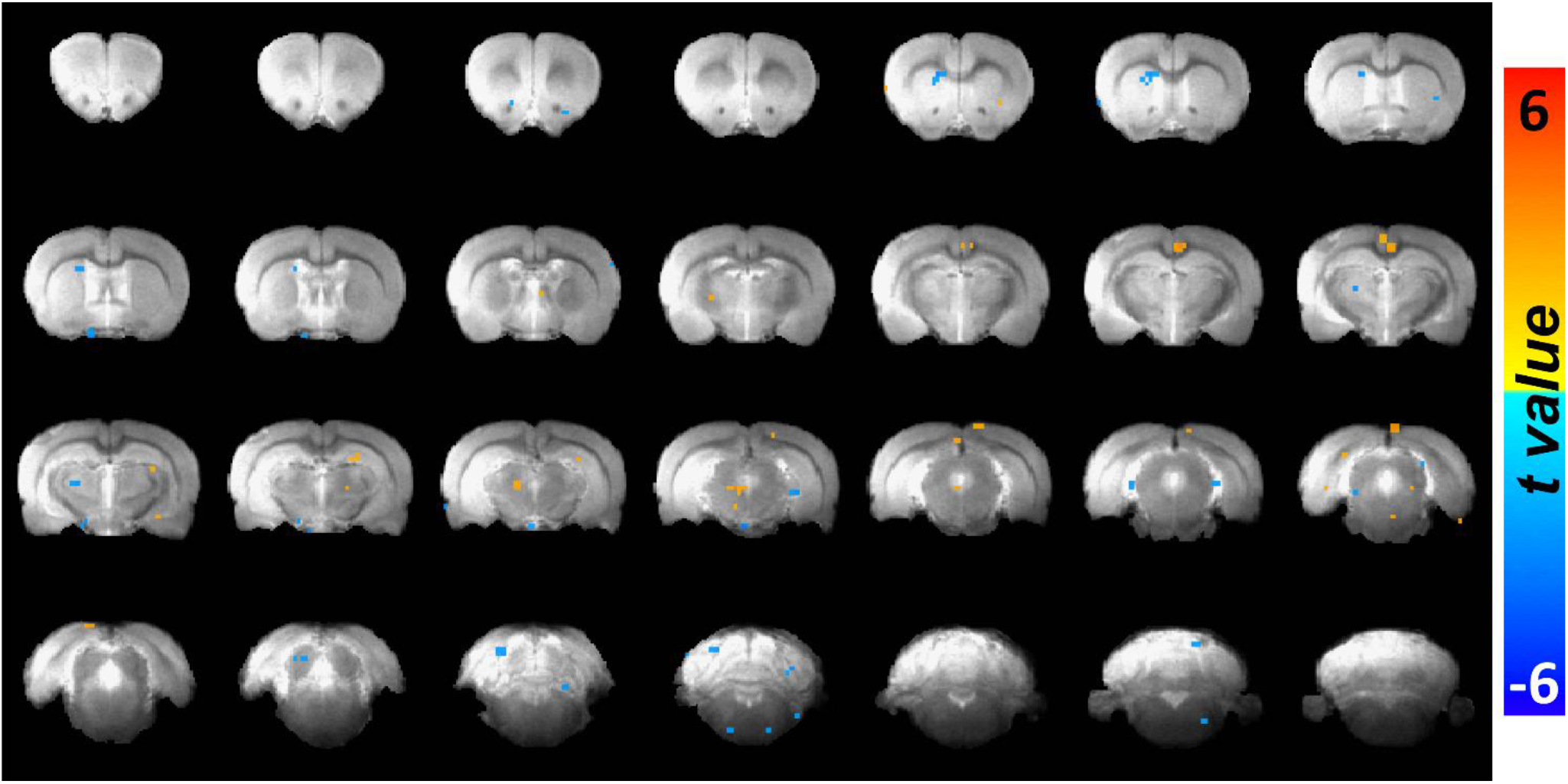
**b)** Averaged correlation maps between ACC 1-4Hz Ca^2+^ power and each voxel of fMRI signals when ACC 1-4Hz Ca^2+^ power and pupil dynamics are not negatively correlated (n=19). The overlay shows voxels with t-value < −2.25 and t-value > 2.25.

**Supplementary Figure 7.**
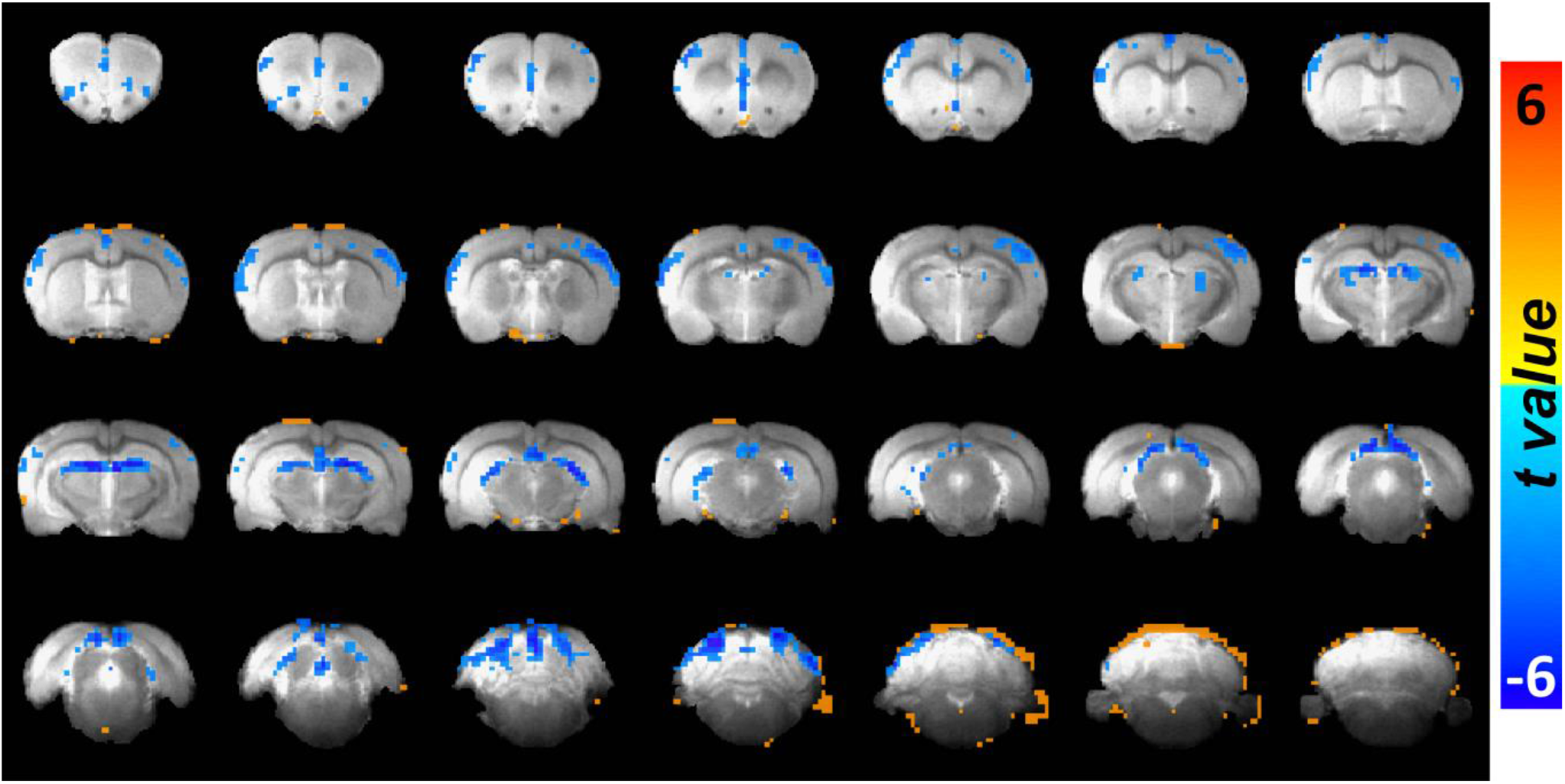
Correlation maps between each fMRI voxel and pupil dynamics from all trials (n=61). The overlay shows voxels with t-value < −2.54 and t-value > 2.54.

**Supplementary Figure 8.**
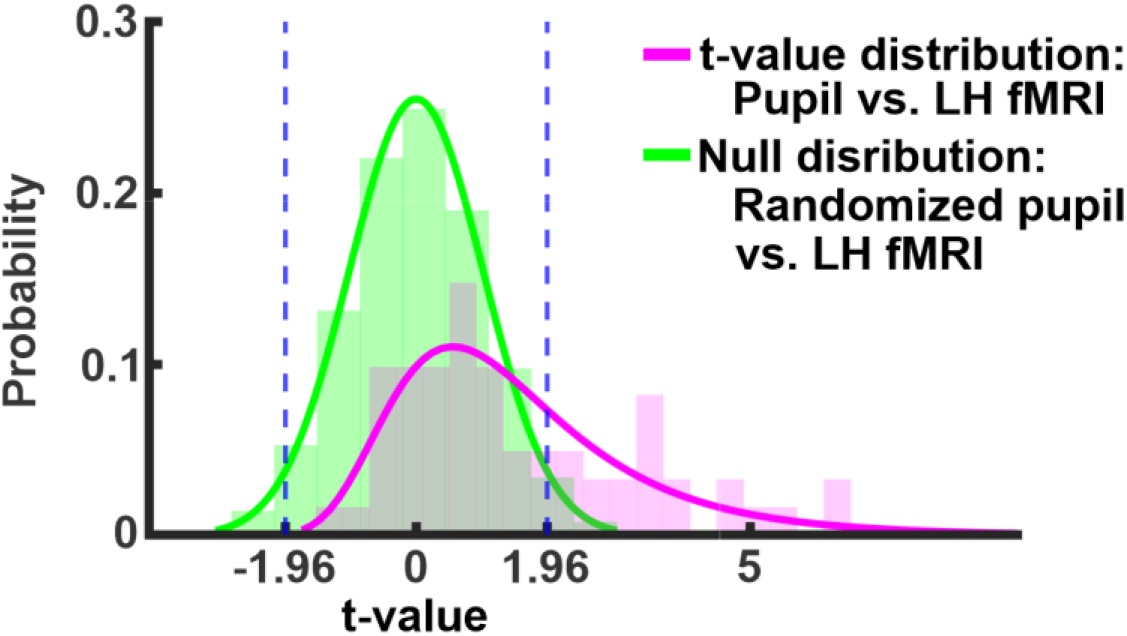
Determination of the threshold for positive and negative correlation between LH fMRI signals and pupil fluctuation. The pupil signals were first randomized, and t-values between this randomized pupil signal and all LH fMRI voxels (AP= −3.2mm) were calculated. Then, permutation was conducted with 10,000 repetitions across 61 trials. The 95% confidence interval of t-values from this permutation test was between −1.96 and 1.96 (indicated with dashed blue line). The distribution of t-values between randomized pupil and LH fMRI is in green and the distribution between pupil and LH fMRI is in pink (same as Fig.4b).

